# Guardians of the cell: The coccosphere prevents bacterial attack in a heavy calcifying coccolithophore

**DOI:** 10.1101/2025.08.12.669917

**Authors:** Sophie T. Zweifel, Richard J. Henshaw, Roberto Pioli, Clara Martínez-Pérez, Uria Alcolombri, Zachary Landry, Roman Stocker

## Abstract

Coccolithophores are responsible for 40–60% of marine calcium carbonate production. This occurs through the biomineralization of extracellular calcium carbonate plates that encase the cell in a structure called the coccosphere. Despite its central role in ocean biogeochemistry, the function of coccolithophore calcification remains unresolved. One hypothesis is that the coccosphere acts as a physical shield, deterring predators and microbes. While its protective role has been investigated against grazers and viruses, its function in bacterial defense remains untested. Here, we investigate the interaction between heavily calcified *Coccolithus braarudii* and the bacterial pathogen *Phaeobacter inhibens*, known for its lethal ‘Jekyll and Hyde’ relationship with the bloom-forming *Gephyrocapsa huxleyi*. We find that in *C. braarudii*, no *P. inhibens* pathogenicity is observed—unless the algae are decalcified. Upon decalcification, the relationship with *P. inhibens* becomes pathogenic, leading to algal cell death. Mortality of decalcified cells is specific to interactions with *P. inhibens* and is attachment-mediated: no toxicity is observed when cells are exposed to *P. inhibens* supernatant or to growth-inhibiting concentrations of indole-3-acetic acid— identified in the *P. inhibens–G. huxleyi* system. Attachment requirement is further supported by scanning electron microscopy, which reveals extensive bacterial colonization on decalcified but not on calcified *C. braarudii* with *P. inhibens*. These findings provide the first experimental evidence that the coccosphere acts as a physical barrier against bacterial attack, underscoring its defensive role in coccolithophores.

## Introduction

Phytoplankton are responsible for an estimated 50% of global primary production^1,2^, with nearly 10% of their collective biomass comprised of a group of calcifying microalgae called coccolithophores^3,4^. Beyond their important role in the organic carbon pump through photosynthesis^5^, coccolithophores mineralize an estimated 40-60% of marine calcium carbonate^3,6,7^, thus playing a major role in the inorganic carbon cycle. This ultimately results in the annual sequestration of over a billion tons of calcium carbonate into the deep ocean when these organisms sink upon cell death^8^.

Coccolithophores mineralize calcium carbonate (CaCO3) into intricate plates known as coccoliths, which are extruded and arranged—typically in one or more interlocking layers—forming a complete or near-complete shell called the coccosphere^9,10^. This mineralized covering makes coccolithophores a key model system for biomineralization^9–14^. Significant work has been dedicated towards understanding the intracellular mechanisms that govern this process^15–17^ and how environmental conditions affect calcification^6,13,18^. Calcification of the cell body can consume as much as 30% of a cell’s total photosynthetic energy budget^9^. Thus, we can expect that the shell must provide a considerable ecological benefit, yet this benefit has remained unclear. Understanding the ‘why’ of calcification is a major open question in microbial oceanography^9^, whose answer will help to better predict how changes to calcification, due to environmental pressures such as ocean acidification, might ultimately influence the fitness of these globally important organisms.

The function of the coccosphere has been explored through a range of hypotheses and experimental studies, revealing that its function likely differs across coccolithophore species^9^. The deep-dwelling coccolithophore *Schyphosphaera apsteinii* mineralizes a higher number of trumpet-like coccoliths in low light conditions: its coccoliths have been proposed to improve light collection^19^. In contrast, shallow-dwelling species, such as *Geopharycapsa huxleyi* (formerly *Emiliania huxleyi*^20^), have flatter coccoliths that can serve as photoprotection^21^. The process of calcification through precipitation of CaCO_3_ has also been shown to shuttle CO_2_ to the immediate surroundings of the cell, which might enhance photosynthesis^22–24^. Other hypotheses have focused on the role of the coccosphere on cell interactions, as a protective barrier against grazing by zooplankton or infection by bacteria or viruses^9^. Although predation rates by dinoflagellates do not differ between calcified, non-calcifying, and artificially decalcified *G. huxleyi* cells^25^, dinoflagellates that consume calcified cells have been found to grow more slowly^26^. A further function of the coccosphere could be as a defensive “shell” or mechanical barrier to protect the cell from attack from viruses and pathogens. Investigation into viral infection of *G. huxleyi* has shown that the coccosphere offers only limited protection against viruses^27^ and in two studies the coccosphere was found to even enhance infection^28,29^. Given that marine viruses can be as small as 20 nm in diameter, while the smallest reported gaps in the *G. huxleyi* coccosphere are 200 nm in size^9,30^, it is expected that the coccosphere only provides limited (if any) protection against viral infection^9^. In contrast, the physical structure of the coccosphere would be almost impenetrable for bacteria, yet the hypothesis that the coccosphere serves as protection against bacteria has to date remained untested.

Here, we investigate the role of the coccosphere in protection from bacterial attack with the coccolithophore *Coccolithus braarudii*: a heavily calcified coccolithophore species and the dominant calcifier in subpolar regions such as the North Atlantic^31,32^. The choice of *C. braarudii* over model organism *G. huxleyi* has two main reasons. Firstly, unlike *G. huxleyi*, which can persist in a non-calcifying diploid state, *C. braarudii* exhibits a strict dependence on its coccosphere—its cell cycle arrests when calcification is inhibited^10,32,33^. Secondly, the extensively characterized “Jekyll-Hyde” interaction between *G. huxleyi* and bacterium *Phaeobacter inhibens*, whereby *P. inhibens* initially releases growth promoting concentrations of indole-3-acetic acid (IAA), but switches to produce both growth-inhibiting concentrations of IAA and algicidal molecules once the algae senesce, occurs regardless of *G. huxleyi*’s calcification state. The *P. inhibens* mediated culture collapse has been reported in both calcifying and non-calcifying *G. huxleyi* strains^34,35^.

In this work, we show that the coccosphere of *C. braarudii* protects the cell from opportunistic bacterial attack. When exposed to coccolithophore-associated bacteria, including *P. inhibens*, artificially-decalcified cultures of diploid *C. braarudii* rapidly collapse, whereas calcified cultures remain unaffected. This response is independent of the method of decalcification. Scanning electron microscopy (SEM) imaging demonstrates that whilst *P. inhibens* attaches to decalcified algae cells, calcified cells appear free from bacterial attachment. This response is shown to be strain-specific, with a panel of bacterial isolates from a previous mesocosm bloom event, *C. braarudii*’s co-isolated consortium, and known coccolithophore-pathogens tested with a range of outcomes from slightly elevating algal growth to full culture collapse. Finally, with *P. inhibens* we show that direct attachment to the decalficied cell is most likely required to induce cell death in *C. braarduii*, with neither co-culture supernatants or exposure to growth-inhibiting concentrations of IAA having a detrimental effect on decalcified cultures. Our results provide the first experimental evidence that the coccosphere can provide a mechanical barrier to opportunistic bacterial attack.

## Materials and Methods

### Algal strains and growth conditions

The xenic diploid strain *Coccolithus braarudii* RCC1200 was grown at 14 °C on a 12:12 diurnal cycle at light intensity of 30 µmol m^-2^ s^-1^ in L1 supplemented synthetic ocean water (SOW, pH = 8)^36^.

### Artificially removing of the coccosphere

For EDTA decalcification, 0.02M EDTA was added to the algal culture for five min^37^, then the cultures were washed and resuspended (1200 relative centrifugal force, RCF, for 5 min) in fresh L1 SOW media. Control (calcified) samples followed identical washing conditions. Cell counts for the decalcified treatments were then adjusted to be commensurate with the washed calcified treatments using flow cytometry (see Methods *Cell Counting*).

For HCl decalcification, 1M HCl was added drop-wise until the culture medium reached pH 2. Cultures were briefly agitated by manual shaking and then neutralized back to pH 8.2 with 1M NaOH^10^. The decalcified and calcified controls were then washed and resuspended (1200 RCF, 5 min) in fresh L1 SOW media. Cell counts for control and treatment conditions were adjusted to the same starting cell densities using flow cytometry (see Methods *Cell Counting*).

### Bacterial growth and addition to algal cultures

Bacterial strains *Marinomonas* 5a1, *Vibrio* 6e7, *Alteromonas* 4a6 and 1d6 were previously isolated from an induced *G. huxleyi* mesocosm bloom in Bergen^38^, DMSZ17395 (*Phaeobacter inhibens*) was obtained from the Leibniz Institute DSMZ-German Collection of Microorganisms, and *Roseovarious nubinhibens* and *Sulfitobacter sp*. D7 was isolated and provided by Assaf Vardi. The *C. braarudii* RCC1200 bacteria isolate and consortium were isolated from the RCC1200 algal culture using 100% Marine Broth medium (Difco, CAS 10043-52-4) agar plates. All strains were preserved in 20% glycerol solution and stored at −80 °C. Bacteria were inoculated in 100% Marine Broth from frozen stocks 24 h prior to use and grown overnight at 30 °C with constant agitation (200 revolutions per minute, RPM). Cultures were collected at stationary phase at optical density (OD) 1.0–1.8 and were washed and resuspended three times (4000 RCF, 5 min) in SOW. Algal cultures were inoculated at stationary phase (14 days old) with the washed bacteria at the specified concentrations (see below *Cell counting*) and were then incubated at 18 °C on a 14:10 diurnal cycle at light intensity 100 µmol m^-2^ s^-1^.

For supernatant experiments (Fig. 3C), a co-culture was prepared from *P. inhibens*, which was grown and washed as specified above; however, in the final washing step the medium was replaced with either *C. braarudii* culture or cell-free *C. braarudii* supernatant rather than SOW. These *P. inhibens* cultures were then incubated for 24 h at 18 °C on a 14:10 diurnal cycle at light intensity 100 µmol m^-2^ s^-1^ before cells were removed (0.2 µm filter, Filtropur) and the resultant filtrate used to inoculate a separate *C. braarudii* culture with the filtered spent medium at a 1:10 filtrate:algae culture volume ratio.

For experiments testing the effect of indole-3-acetic acid (Figs 3B, S6) (IAA), IAA was prepared fresh before each experiment in L1 SOW medium with filter sterilization (0.2 µm filter, Filtropur). IAA was first dissolved in 50% ethanol prepared with L1 medium to ensure solubility, then diluted into the algal culture to a final ethanol concentration of ≤ 0.1%. This low concentration is well below the 1% threshold previously shown to have no effect on algal growth^39^.

### Microscopy and cell counting

*C. braarudii* cells in four different treatment conditions (either calcified or decalcified, and with/without addition of *P. inhibens*) were gently pipetted into a six-channel IBIDI chip (μ-Slide uncoated VI 0.4, Cat.No. 80601) and the ports sealed with strips of parafilm. The chip was imaged using phase-contrast microscopy (Nikon Ti-E; 10× 0.3 NA objective), with an image taken once per hour for a period of 100 hours using a CMOS camera (Orca-Flash 4.0 Hamamatsu), and cells counted manually.

Flow cytometry was used to count both algal and bacterial cells using a cytometer (Beckman Coulter, CytoFLEX S) equipped with a 488 nm laser. For algal cell counts, forward scatter (FSC) and red autofluorescence (filter for Peridinin–chlorophyll protein complex PerCP-A) were recorded. For bacterial counts, separate aliquots were stained with SYBR Green (5 µM final concentration, Sigma Aldrich CAS 163795-75-3) and incubated for 10 min in the dark. Cell counts were then obtained by recording FSC and green fluorescence (filter for fluorescein, FITC).

Bacterial counts were additionally obtained as counts of colony forming units (CFUs) after plating. Aliquots of 100 µL sequentially diluted culture were spread onto 100% Marine Broth Agar (Difco, CAS 90000-518) and incubated at 30 °C for 48 h before counting. Among samples at different dilutions, plates with ~100 colonies were counted. In quantifying CFUs, *P. inhibens* was distinguishable from other bacteria in the culture due to their formation of brown colonies after 48 h (SI Figure S3).

### Scanning electron microscopy

200 µL samples of *C. braarudii* liquid cultures were fixed with 1% (w/v) glutaraldehyde for 1 h then gently pipetted onto 0.01% poly-L-lysine coated hydrophilized silicon wafers. The wafers were then incubated at ambient temperature for 10 min, then sequentially immersed in 2.5% glutaraldehyde solution (SOW, 27 practical salinity units, PSU), 1% osmium tetroxide, and SOW, for 5 min each. Next the wafers underwent an ethanol drying series consisting of 2 min immersion in 0%, 30%, 50%, 70%, 90%, and 100% ethanol, followed by three times water-free 100% ethanol. Critical point dryer (CPD 931 Tousimis, ETH Zurich Microscope Facility ScopeM) with a cell monolayer protocol was then used and the wafers adhered to aluminum stubs with silver paint. After 24 h degassing, the wafers were sputter coated with 4 nm platinum palladium (CCU-010 Metal Sputter Coater Safematic, ETH Zurich Microscope Facility ScopeM). Cells were imaged using an extreme high resolution (XHS) TFS Magellan 400 microscope (50 pA current, 5 kV accelerating voltage, ETH Zurich Microscope Facility ScopeM). Backscattered electrons were imaged with a CBS detector and secondary electrons with a TLD detector, both using immersion.

### Statistics and reproducibility

Graphpad Prism 10.4.1 (GraphPad Software, La Jolla, CA, USA) was used for all statistical analyses and graphing. Data are all reported as the mean ± standard error of the mean (SEM). In all cases, a *p*-value of *p* < 0.05 was considered statistically significant.

## Results

Here, we demonstrate that the coccosphere of calcified, diploid *C. braarudii* cells acts as an effective barrier against opportunistic bacterial attack. Calcified *C. braarudii* cultures are unaffected when exposed to the known *G. huxleyi* pathogen *Phaeobacter inhibens*^34,35,40^ across a wide range of initial bacterial concentrations. In contrast, artificially decalcified *C. braarudii* cultures exhibit reduced growth at low bacterial concentrations and completely collapse due to bacteria-induced cell death at high bacterial concentrations. We observed this collapse of decalcified *C. braarudii* cultures uniquely with *P. inhibens* (and to a lesser degree with a *G. huxleyi* associated bacterium, *Vibrio* 6e7) and not with a range of other coccolithophore-associated bacteria. Furthermore, we found growth inhibition could not be triggered by addition of high concentrations of indole-3-acetic acid (IAA, 1000 uM) as has been seen with *G. huxleyi*^34,39,40^, where it has been suggested that *P. inhibens* switches from a mutualistic relationship to a pathogenic one partially by up-regulating IAA. The triggered killing of decalcified *C. braarudii* by *P. inhibens* is shown to require direct attachment, as no cell collapse was observed when cultures were exposed to *P. inhibens* supernatant alone. This conclusion is further supported by scanning electron microscopy, which reveals substantial bacterial attachment only on decalcified cells exposed to *P. inhibens*, while exposed calcified cells remain largely free from bacterial attachment. These findings demonstrate that the coccosphere of *C. braarudii* functions as a protective barrier to prevent opportunistic attack by pathogenic bacteria, expanding the known functions of the coccosphere in coccolithophores and providing a blueprint for similar investigations in other coccolithophore species.

### Calcification protects Coccolithus braarudii from Phaeobacter inhibens–mediated cell death

In order to investigate the role of *C. braarudii’*s coccosphere, we first investigated methods to decalcify *C. braarudii* without lasting effects on either the cells’ growth or their ability to recalcify. We found that treatment with ethylenediaminetetraacetic acid (EDTA, 0.02M EDTA, 5 min, see Materials and Methods) efficiently decalcified the cells. This was confirmed both visually (SI Video S1) and via flow cytometry, with decalcified cells producing reduced forward scatter (FSC-A) and elevated fluorescence signals when compared to their calcified counterparts (SI Fig. S1). We found 95% of the population to decalcify upon EDTA treatment, compared to only 4% in no-EDTA controls (Fig. 1A). Post-treatment, flow cytometry tracking and time-lapse microscopy revealed that cells began recuperating quickly, with recalcification followed by cell growth progressing in parallel with untreated controls and reaching comparable levels by day 10 (Fig. 1A, Fig. 1B). Timelapse microscopy and scanning electron microscopy confirmed the emergence of visible coccoliths within three days post-decalcification (Fig. 1BCD) and 90% of the population recalcified by day 10 post-treatment (Fig. 1A). Similar decalcification and recovery results were also achieved using HCl treatment (1 M HCl for 5 minutes at pH 2, followed by neutralization to pH 8; SI Fig. S2). We chose to use the EDTA treatment for decalcification in the rest of this work, as it resulted in higher cell growth compared to the HCl treatment (SI Fig. S2) and has been shown to be less likely to damage cellular structures and interfere with physiological processes^41,42^.

**Fig. 1.**
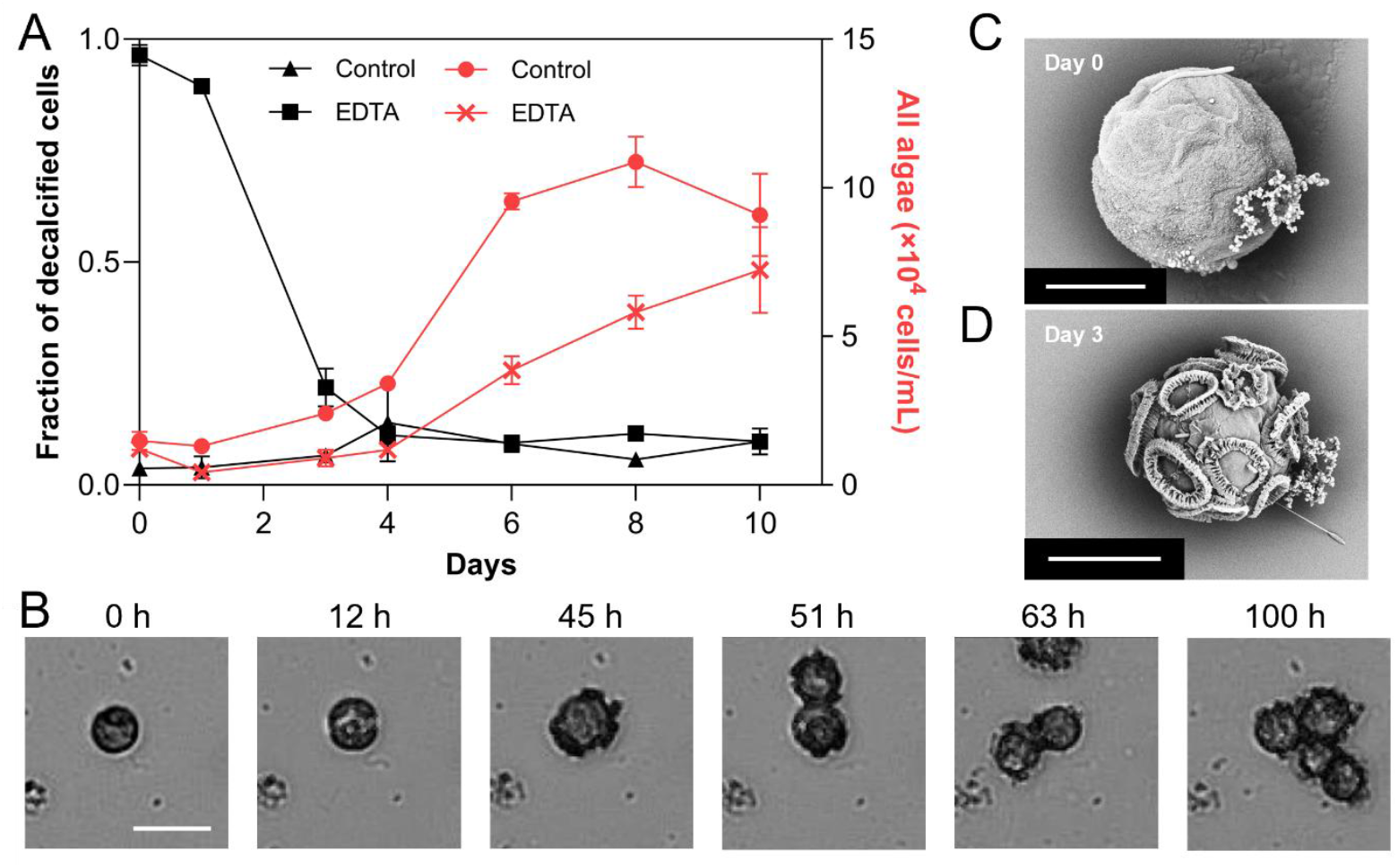
Tracking and imaging of artificially decalcified *C. braarudii* cells. *(A)* Cell counts (red, right axis) and decalcified fraction (black, left axis) as a function of time for untreated cultures (red circles and black triangles) and cultures decalcified using EDTA (black squares, red crosses). Note the recalcification and cell growth in the EDTA treatment over the course of ten days. Symbols represent the mean with SE bars for *n* = 3 biological replicates. *(B)* Timelapse of EDTA-decalcified cells over the course of 100 hours, showing recalcification and growth (scale bar = 10 µm). *(C,D)* SEM images of a cell (*C*) immediately after decalcification with EDTA and (*D*) three days after decalcification (scale bar = 5 µm).

The marine bacterium *Phaeobacter inhibens* exhibits a well-characterized “Jekyll-and-Hyde” interaction with the coccolithophore *Geopharycapsa huxleyi*^34,35,39,40^, wherein *P. inhibens* initially promotes algal growth, leading to elevated cell densities relative to unexposed algae cultures. This mutualistic phase is transient, however: upon sensing specific algal-derived signals, *P. inhibens* switches to a pathogenic state, producing bioactive secondary metabolites that induce rapid algal mortality and culture collapse both in calcifying and non-calcifying *G. huxleyi* strains^34,35^. This led us to investigate whether the same interaction occurs between *P. inhibens* and *C. braarudii* cultures along with its co-isolated bacterial community. We combined *C. braarudii*, both calcified and EDTA-decalcified, with different initial concentrations of *P. inhibens* (10^4^ to 10^6^ cells/mL) and monitored the algal population through both light microscopy and flow cytometry. These bacterial concentrations—particularly 10^6^ cells/mL—are on the higher end of what is observed in natural seawater, but fall within the range reported for dense microbial aggregations such as those occurring in phytoplankton blooms, where localized bacterial abundances can exceed 10^7^ cells/mL^43,44^. We selected this value to ensure a sufficiently strong and measurable interaction between host and bacterium across treatments, while acknowledging that the upper range likely represents only limited environmental conditions.

The growth of calcified *C. braarudii* was unaffected by exposure to *P. inhibens* at either 10^5^ or 10^6^ cells/mL over a 10-day period, as confirmed by both time-lapse microscopy and flow cytometry (Fig. 2A–B). In stark contrast, EDTA-decalcified *C. braarudii* showed marked sensitivity to *P. inhibens* exposure: at 10^5^ cells/mL, cell numbers began to decline after two days, while exposure to 10^6^ cells/mL resulted in rapid population collapse within 15 hours (Fig. 2C). Flow cytometry further confirmed this dose-dependent response, revealing a significant reduction in algal cell numbers for the 10^5^ and 10^6^ treatments by days 7–8 (Fig. 2D). In comparison, cultures exposed to 10^4^ *P. inhibens* cells/mL showed no significant difference from untreated controls over our experimental time of 8 days. These results demonstrate that decalcified *C. braarudii* cells are highly susceptible to *P. inhibens*, but only above a certain bacterial threshold. Notably, this vulnerability was independent of the decalcification method, with similar results observed using HCl treatment (SI Fig. S2).

**Fig. 2.**
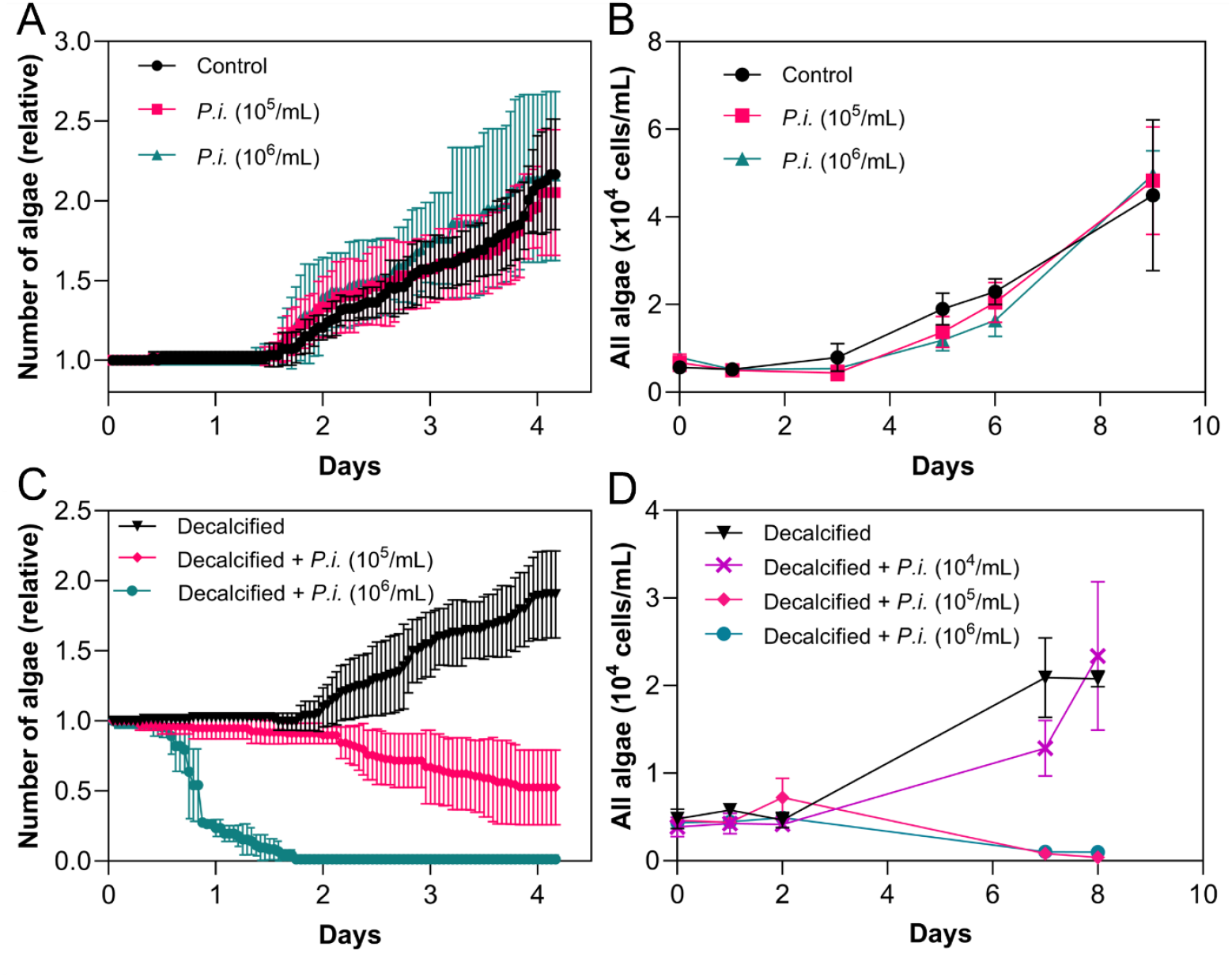
Differential response of calcified (A, B) and EDTA-decalcified (C,D) *C. braarudii* to *P. inhibens* (‘Pi’) bacteria, assessed via time-lapse microscopy *(A, C)* and flow cytometry *(B, D)*. Cell numbers in the microscopy measurements (*A, C*) were normalized to those at time 0. *(A)* Single-cell microscopy of calcified *C. braarudii* reveals no significant difference in growth between untreated controls (black circles) and cultures inoculated with *P. inhibens* at 105 (pink squares) or 106 cells/mL (green triangles) (two-way ANOVA, F2,12 = 0.48, *p* = 0.63; n = 5). *(B)* Flow cytometry measurements further confirm no difference in growth between the same three treatment (two-way ANOVA, F_1,4_ = 0.71, p = 0.45; n = 3). *(C)* Single-cell microscopy of EDTA-decalcified *C. braarudii* reveals a marked decrease in cell numbers, compared to the control (black triangles), upon addition of *P. inhibens* at 105 cells/mL (green circles) and 106 cells/mL (two-way ANOVA, F_2,12_ = 134.4, p < 0.0001; Dunnett’s test, p < 0.05; n = 5). *(D)* Flow cytometry measurements show a dose-dependent decline in decalcified populations exposed to *P. inhibens* at 104 (purple crosses), 105 (pink diamonds), and 106 (green circles) cells/mL. The reduction in algal cell number compared to the control was significant for 105 and 106 cells/mL on day 7 (106 Pi: p = 0.0310; 105 Pi: p = 0.0311) and day 8 (106 Pi: p = 0.001; 105 Pi: p = 0.0003) (two-way ANOVA, F_3,8_ = 104.7, p < 0.0001; one-tailed Dunnett’s test; n = 3 biological replicates).

**Fig. 3.**
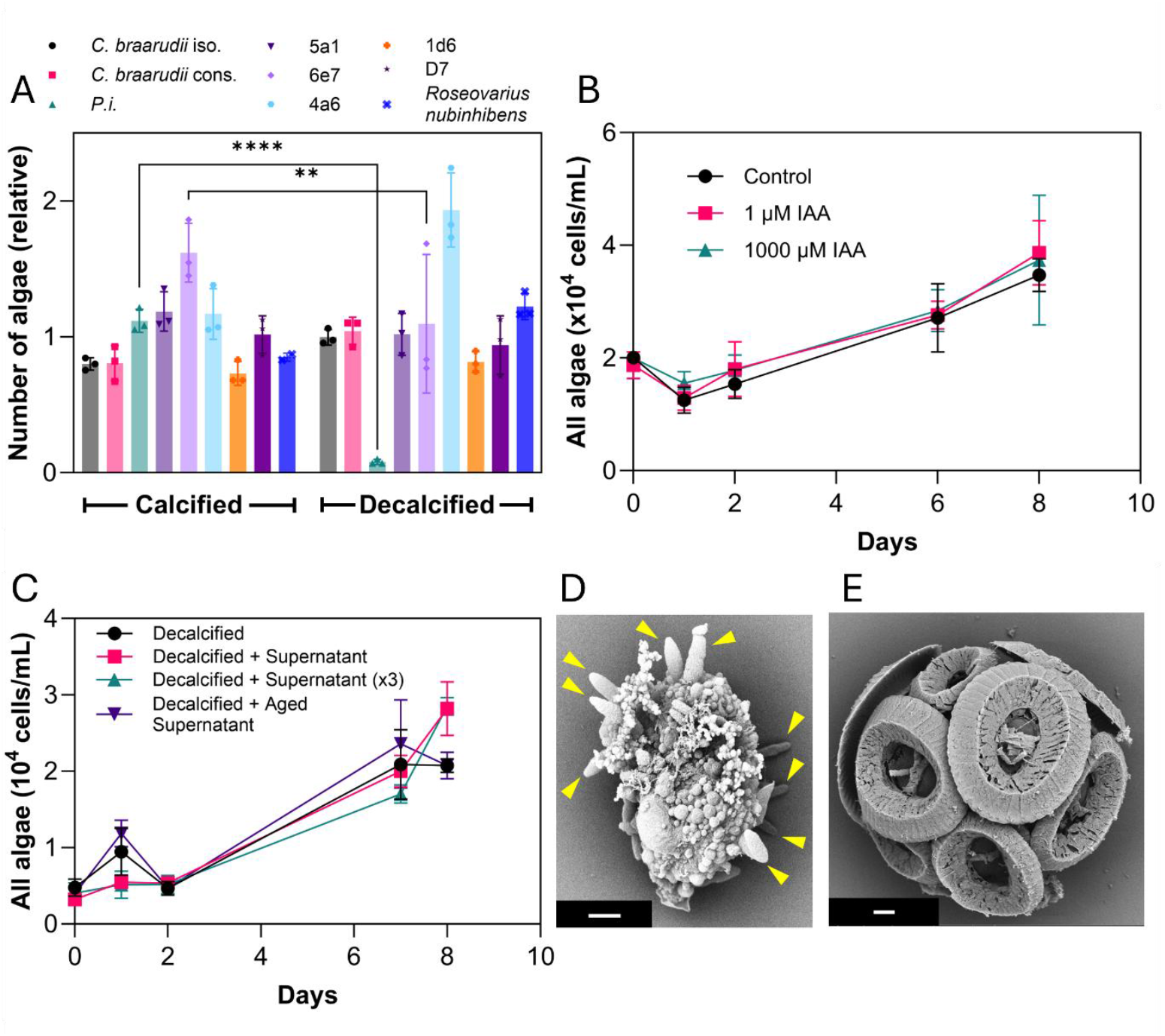
The collapse of decalcified *C. braarudii* cultures upon bacterial exposure is bacterial-strain-specific and requires physical attachment. *(A)* Normalized concentrations of calcified and EDTA-decalcified *C. braarudii* after 8 days of exposure to different coccolithophore-associated bacterial strains (different symbols and colors). Cell counts were obtained by flow cytometry and are normalized first to day 0 values and then to the mean cell count of the respective untreated controls (calcified or decalcified) on day 8. Most bacteria had no significant impact, except for *P. inhibens* DSMZ17395 and *Vibrio* strain 6e7, which triggered significant decrease in algal cell counts in decalcified but not in calcified cultures (one-way ANOVA, F8,18 = 12.01, p < 0.0001; Šídák’s post hoc test; **p < 0.01, ****p < 0.0001). *(B)* Time-resolved cell counts obtained by flow cytometry for decalcified *C. braarudii* exposed to indole-3-acetic acid (IAA) at concentrations previously reported to promote growth (1 µM) or have algicidal effects (1000 µM) in *G. huxleyi*^34^. No significant effect on cell growth of IAA at either concentration was observed (two-way ANOVA, F2,6 = 0.95, p = 0.44; n = 3). *(C)* Time-course analysis of decalcified *C. braarudii* populations exposed to various *P. inhibens* cell-free supernatants: (i) a single dose from *P. inhibens* pre-incubated in *C. braarudii* spent medium, (ii) three doses of the same, delivered on days 0, 1, and 2, and (iii) the supernatant from a 7-day-old collapsed co-culture. None of these treatments led to a significant change in cell numbers (two-way ANOVA, F3,6 = 0.52, p = 0.68; n = 3), suggesting that direct contact is necessary to induce collapse. *(D, E)* Scanning electron micrographs of decalcified *C. braarudii* cells three days post-decalcification. Yellow triangles (*D*) denote bacteria attached to an EDTA decalcified *C. braarudii* following exposure to 10^6^ cells/mL of *P. inhibens. (E)* No attachment was observed on calcified *C. braarudii* cells inoculated with the same concentration of *P. inhibens*. Scale bars = 1 µm.

Exposure of *C. braarudii* to *P. inhibens* was associated with an increase in the other bacterial populations present in the algal cultures. Bacteria were quantified as colony forming units (CFUs, SI Fig. S3), with *P. inhibens* distinguishable from other bacteria by their formation of brown colonies (SI Fig. S4). In both calcified and decalcified treatments, addition of *P. inhibens* resulted in an increase in other bacterial populations (SI Fig. S3A). When inoculated in identical conditions (starting bacterial and algal cell densities), *P. inhibens* exhibited markedly faster growth dynamics when co-cultured with calcified *C. braarudii* cells compared to artificially decalcified *C, braarudii* cells. In the calcified condition, *P. inhibens* grew by more than 22-fold, reaching a peak of 3.7 × 10^7^cells/mL. In contrast, growth in the calcified treatment was much more limited, with only a 5.6-fold increase to a maximum of 8.3 × 10^6^ cells/mL—a reduction of ~345% in peak abundance compared to the decalcified treatment. This drastic increase in bacterial growth in the decalcified treatment is even more notable since decalcified *C. braarudii* exhibit slightly reduced growth relative to their calcified counterparts (Fig. 1A). A sharp drop in *P. inhibens* abundance occurred in the decalcified treatment after day 7, coinciding with the collapse of the algal population (SI Fig. S3A), suggesting a tight coupling between bacterial and host dynamics in the absence of the coccosphere. This concurrent decrease in concentration likely reflects the exhaustion of algal-derived nutrient sources for *P. inhibens*, suggesting that the pathogenic phase is self-limiting in the absence of a sustained host.

### Strain-specificity and mechanisms underlying the collapse of decalcified C. braarudii cultures exposed to P. inhibens

To determine whether the collapse of decalcified *C. braarudii* cultures is specific to *P. inhibens*, we tested eight additional bacterial strains on artificially decalcified cultures (Fig. 3A). These were: (i) a *C. braarudii*-associated bacterial consortium derived from long-term laboratory cultures, (ii) a single isolate from the long-term laboratory consortium, (iii) four isolates obtained from a *G. huxleyi*-stimulated mesocosm bloom event in Bergen^38^ (5a1, 6e7, 4a6, 1d6), and (iv) two other known *G. huxleyi* pathogens *Sulfitobacter sp*. D7 and *Roseovarius nubinhibens*^45,46^. Among all treatments, only *P. inhibens* and one specific mesocosm isolate, *Vibrio* 6e7, caused a significant decrease in *C. braarudii* cell numbers in decalcified cultures (Fig. 3A). In contrast to *P. inhibens*, the other known *G. huxleyi* pathogens *Sulfitobacter* D7 and *R. nubinhibens* had no effect on *C. braarudii* growth in either calcified or decalcified cultures. Moreover, exposure to both *Alteromonas* 4a6 and the *C. braarudii* consortium, resulted in slightly elevated *C. braarudii* cell numbers in decalcified cultures compared to calcified controls. These findings demonstrate that the bacteria-induced collapse of decalcified *C. braarudii* is highly specific, with different bacterial isolates inducing varying effects on the algal cultures.

To investigate the mechanism underlying the collapse of decalcified *C. braarudii* cultures when exposed to *P. inhibens*, we first tested whether the same algicidal phenomena present in *G. huxleyi–P. inhibens* system was also responsible for the cell death of *C. braarudii*. With *G. huxleyi–P. inhibens* systems, the transition of *P. inhibens* to a pathogenic state is marked by increased production of indole-3-acetic acid (IAA), a growth hormone that—at elevated concentrations—is sufficient to induce algal culture collapse without direct cell-cell contact or attachment^34,35,40^. Here, we inoculated calcified and decalcified *C. braarudii* cultures with IAA at two concentrations, 1 µM and 1000 µM, previously shown to be respectively growth-promoting and growth-inhibiting to *G. huxleyi*^34,39,40^. No significant effect on *C. braarudii* population growth was observed at either concentration with either decalcified (Fig. 3B) or calcified (SI Fig. S5) cultures. These findings indicate that IAA-mediated growth inhibition is not contributing to the observed collapse in artificially decalcified *C. braarudii*. Instead, the interaction is likely driven by other bacterial metabolites and/or requires direct cell–cell contact—highlighting a distinct, contact-dependent mechanism of pathogenicity.

To investigate whether direct attachment is necessary for *Phaeobacter inhibens* to induce cell death in decalcified *C. braarudii*, we tested whether bacterial exudates alone could trigger the same collapse phenotype. We designed three treatments utilising the supernatants of co-cultures: (i) Single supernatant addition: *P. inhibens* was grown overnight, washed, and then incubated for 24 h in cell-free spent medium from a *C. braarudii* culture. After this incubation, bacterial cells were removed by filtration and the resulting supernatant was added once (day 0) to a fresh decalcified *C. braarudii* culture. (ii) Repeated supernatant additions: The same supernatant as in point (i) was added on three consecutive days (days 0, 1, and 2) to test whether cumulative exposure or higher metabolite concentration would induce algal cell death. (iii) Post-collapse co-culture supernatant: Supernatant was collected from a 7-day-old co-culture of decalcified *C. braarudii* and *P. inhibens* that had already undergone collapse. After removing all cells, this mixture of metabolites was added to a fresh decalcified *C. braarudii* culture to test whether toxic compounds accumulated during collapse could trigger algal cell death without direct bacterial contact. Across all three treatments, no significant changes in algal cell growth were observed over an 8-day period compared to untreated decalcified controls (Fig. 3C). These results indicate that soluble exudates from *P. inhibens* alone are insufficient to induce cell death in *C. braarudii* and that direct attachment is likely required.

The need for attachment is further supported by scanning electron microscopy measurements of the co-cultures, which showed substantial bacterial attachment to decalcified *C. braarudii* treated with *P. inhibens* (Fig. 3D), whereas no attachment was observed on calcified cells. No attachment was observed on decalcified cells not treated with *P. inhibens* either (Fig. 3E, 1C), indicating that the attached bacteria in the *P. inhibens*-treated cultures are likely *P. inhibens*, as opposed to members of the natural consortium of *C. braarudii* (it is also possible that the presence of *P. inhibens* triggers the attachment of other members of the xenic culture)Together, these findings indicate that the pathogenic behavior of *P. inhibens* is likely contact-dependent and ultimately that the coccosphere of *C. braarudii* forms a mechanical barrier preventing opportunistic attack by bacteria that would otherwise result in cell death. This result supports the hypothesis that calcification may shield some phytoplankton from potentially harmful bacterial interactions.

## Discussion

Among multiple hypotheses on coccosphere functionality, one proposes that the tightly interlocked coccoliths could provide a mechanical barrier against bacterial attack^9^ by reducing or preventing bacterial attachment and penetration. Previous observations of the attachment of *P. inhibens* to only naturally decalcified *G. huxleyi*^34^, combined with reports of the calcification ability of *G. huxelyi* becoming lost over many generations when maintained in the laboratory^10,47,48^, which are frequently kept under axenic conditions, suggest that bacteria could influence the algal cell investment in calcification. Here, we provide direct experimental evidence that the coccosphere of a heavily calcifying species, *C. braarudii*, offers physical protection against bacterial-induced killing. Upon artificial decalcification (Methods), *C. braarudii* cells become highly susceptible to bacterial attack from specific bacterial strains, including the well-characterized *G. huxelyi* pathogen *P. inhibens*. Inoculating artificially decalcified *C. braarudii* cultures with concentrations of *P. inhibens* above 10^5^ cells/mL triggers rapid culture collapse (Fig. 2CD), whereas it has no measurable effect on calcified cultures (Fig. 2AB) despite a naturally occurring decalcified population fraction of approximately 4% (Fig. 1A). A similar result was observed with inoculation with a second bacterial species (*Vibrio* 6e7), isolated from an induced mesocosm bloom^38^. In contrast, no response was observed with five other coccolithophore-associated bacterial strains, highlighting that the response is species-specific. It is important to note that, whilst we demonstrate how *C. braarudii* recovers and eventually re-calcifies after artificial decalcification (Fig. 1), it is possible that stress from the decalcification process increases the susceptibility of the algae to bacterial pathogens. However, any such increase in susceptibility is likely minor, as decalcified *C. braarudii* cultures eventually recover over time, and furthermore do not show any culture collapse when exposed to *P. inhibens* at initial concentrations of 10^4^ cells/mL (Fig. 2D).

This algicidal interaction is likely to have a chemical component—for example, through the production of algicidal compounds, as seen in *P. inhibens–G. huxleyi* system^34,35^. Despite providing significant coverage of the algal cell (Fig. 3D), the coccosphere is not free of gaps and not completely impermeable to small molecules. Thus, diffusible algicidal compounds could reach the cell regardless of calcification state: this process would be favored by bacteria establishing close contact with the algae. However, we observed no negative effect of *P. inhibens* on calcified *C. braarudii* cultures, suggesting that mere proximity to the cell itself is insufficient and that direct physical attachment may be required—perhaps to enable close-range delivery of algicidal compounds or even direct injection into the host cell. This hypothesis is supported by our exudate experiments: exposing both calcified and decalcified *C. braarudii* cultures to exudates from *P. inhibens* did not induce significant cell death (Fig. 3C), nor did treatment with the phytohormone indole-3-acetic acid (IAA; Fig. 3B, SI Fig. S5)—a compound produced by *P. inhibens* and shown to be growth-inhibiting to *G. huxleyi* at high concentrations (1000 µM) without requiring direct contact^34,39,40^. The lack of any detectable effect, even at high IAA concentrations and with exudates, supports the idea that killing in the *P. inhibens– C. braarudii* system is not mediated by diffusible compounds alone, but rather requires direct cell–cell attachment—an interaction that may be physically obstructed by the coccosphere. Notably, the specific compound(s) responsible for inducing algal death in decalcified *C. braarudii* cultures remain unidentified. Whether the same algicidal compounds implicated in *G. huxleyi* mortality are also active here^34,35,39,40^, or whether an alternative mechanism is involved, remains an open question. Here we note that the strikingly different interaction between *C. braarudii* and *P. inhibens* when compared with *G. huxelyi* highlights a functionally distinct and previously uncharacterized alga–bacterium pairing. This system offers a promising new model for disentangling species-specific microbial pathogenesis and the evolutionary diversification of algal defenses.

Our results support the interpretation of the coccosphere functionality for the heavily calcified *C. braarudii* as a defensive “shell” physically protecting the cell body from opportunistic bacterial attachment and subsequent induced cell death. We show that *P. inhibens* proliferated substantially more in decalcified cultures than in calcified ones (peak bacterial populations of 3.7 × 10^7^cells/mL and 8.3 × 10^6^ cells/mL, respectively). In the decalcified algal cultures, the *P. inhibens* population sharply declined after day 7, concurrent with the collapse of the algae culture (SI Fig. S3A). SEM imaging showed that *P. inhibens* attached readily to decalcified cells but not to calcified ones (Fig. 3DE), demonstrating that the coccosphere of *C. braarudii* prevents *P. inhibens* attachment. Prior genomic studies have shown that *P. inhibens* encodes structural genes for a Type VI secretion system (T6SS), which in other species (e.g., *Campylobacter jejuni*) delivers toxic effectors directly into target cells through physical contact using a tube-like structure^49,50^, and can be impaired by extracellular barriers such as capsules or matrices^51^. While T6SS functionality in *P. inhibens* DSM 17395 has not been shown, if active, it could explain the pathogenic effect on naked cells and be neutralized by the presence of the coccosphere. Furthermore, the individual coccoliths forming the coccosphere are coated with coccolith-associated polysaccharides (CAPs) which forms a dense, negatively charged barrier^52,53^ that could also disrupt molecular cues required for bacterial attachment. *P. inhibens* forms polar N-acetylglucosamine-rich adhesins to establish surface contact and form rosettes^54,55^: interference with these structures could prevent direct attachment required for pathogenesis.

For the heavily calcifying coccolithophore *C. braarudii*, our findings demonstrate that at least part of the coccosphere functionality is as a mechanical barrier to protect the cells against opportunistic bacterial attack. This adds to growing evidence that structural armoring in phytoplankton may have appeared, in part, as a defense against biotic threats—not only grazers but also microbial pathogens—mirroring trends observed in other lineages of mineralized or spiny phytoplankton^56–58^. We propose this is achieved by forming a biochemically selective interface which disrupts both bacterial attachment and contact-mediated effector delivery. This dual function highlights the broader ecological role of calcification in modulating host–microbe interactions. Environmental stressors that compromise coccosphere integrity could significantly increase coccolithophore vulnerability, altering bloom dynamics and disrupting the intricate balance between algae, bacteria, and viruses—interactions that ultimately drive large-scale ecological events and nutrient fluxes across marine ecosystems.

## Acknowledgments

We thank Assaf Vardi, Assaf Gal, Oliver Mueller, Johannes Keegstra, Ariel Chazan, Frédéric de Schaetzen, Dieter Baumgartner and Isobel Short for helpful discussions, Russell Naisbit for editing and feedback, and Ela Burmeister for help with experimental work. We thank Assaf Vardi for gifting of strains *Sulfitobacter sp*. D7 and *Roseovarious nubinhibens*. The authors gratefully acknowledge Stephan Handschin, Miriam Lucas, and Anne Greet Bittermann of ScopeM at ETH Zurich for their assistance in electron microscopy imaging.

## Competing Interest Statement

R.S is the co-founder and chief scientific officer of PhAST Crop., a startup working on antibiotic resistance diagnostics. There is no relation between PhAST and this publication.

All other authors declare no competing interests.

## Funding

We gratefully acknowledge funding from a Gordon and Betty Moore Foundation Symbiosis in Aquatic Systems Initiative Investigator Award (GBMF9197; https://doi.org/10.37807/GBMF9197), the Simons Foundation through the Principles of Microbial Ecosystems (PriME) collaboration (grant 542395FY22), Swiss National Science Foundation grant 205321_207488, and the Swiss National Science Foundation Sinergia grant CRSII5-186422, all to R.S. And the HFSP fellowship (LT000192/2018) to ZL.

## Data and Code Availability

Data files used during the current study are publicly available at the ETH Research Collection, which can be accessed at this DOI: https://doi.org/10.3929/ethz-b-000745131.

## Notes

https://doi.org/10.3929/ethz-b-000745131

