## Supplementary material for "Guardians of the cell: The coccosphere prevents bacterial attack in a heavy calcifying coccolithophore": SI Material

Supplementary Figure 1. **Flow cytometry gating for the identification of calcified and decalcified cells in *C. braarudii* population.**

Supplementary Figure 2. **Comparison of decalcification methods for *Coccolithus braarudii***

Supplementary Figure 3. **Counts of *Phaeobacter inhibens* and other bacteria when inoculated into calcified and decalcified *Coccolithus braarudii* cultures.**

Supplementary Figure 4. **Comparison of bacteria colonies formed with different *Coccolithus braarudii* treatments.**

Supplementary Figure 5. **Cell growth response of calcified *C. braarudii* cultures to addition of indole-3-acetic acid.**

Supplementary Video S1. ***Coccolithus braarudii* recalcification and cell division.**

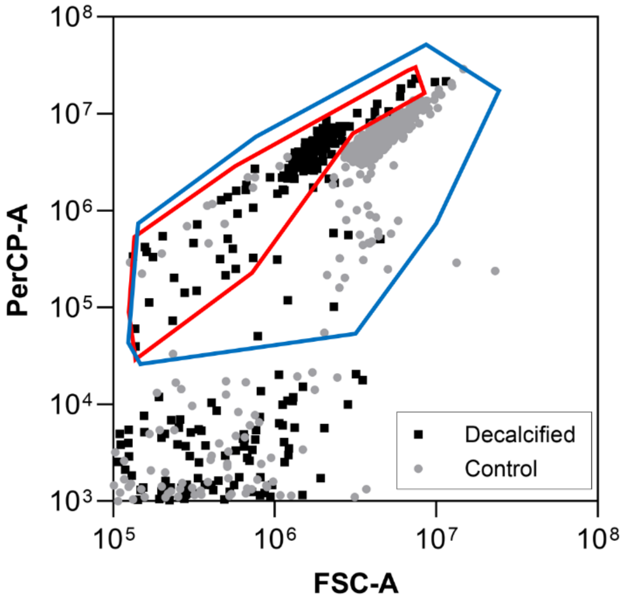


**Supplementary Figure 1. Flow cytometry gating for the identification of calcified and decalcified cells in *C. braarudii* population.** Flow cytometry shows distinct clustering between decalcified (black squares) and control (grey circles) treatments. Populations were identified based on chlorophyll autofluorescence (PerCP-A), with decalcified and control (calcified) cells occupying separate regions of the plot, indicating a shift in cellular optical properties following decalcification with EDTA.

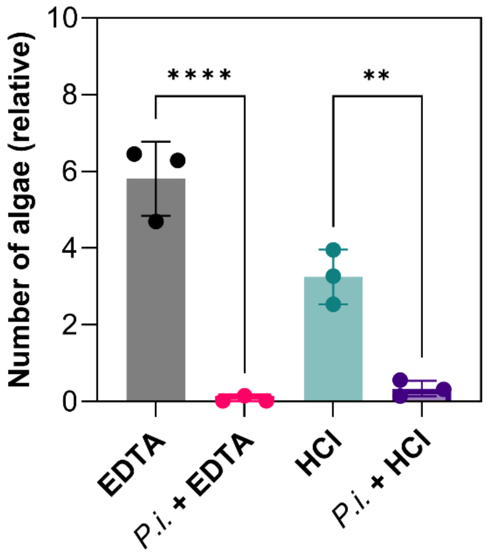


**Supplementary Figure 2. Comparison of decalcification methods in *Coccolithus braarudii* and their effects on susceptibility to *Phaeobacter inhibens*.** *C. braarudii* cultures were decalcified with either HCl (1 M) or EDTA (0.5 M). For each treatment, control conditions received no treatment whereas +*P.i.* (*P. inhibens*) treatments were inoculated with bacteria ($10^6$ cells/mL). Bars show the mean with SE bars for $n = 3$ biological replicates (shown as individual points). Comparison of treatments, two-way ANOVA ($F_{3,6} = 66.78$, $p < 0.0001$). Both *P. inhibens* treatments differ significantly from their respective control (two-tailed Tukey's multiple comparisons test, EDTA $p < 0.0001$, HCl $p = 0.0033$). Algal concentration was normalized to the day zero counts for each treatment group.

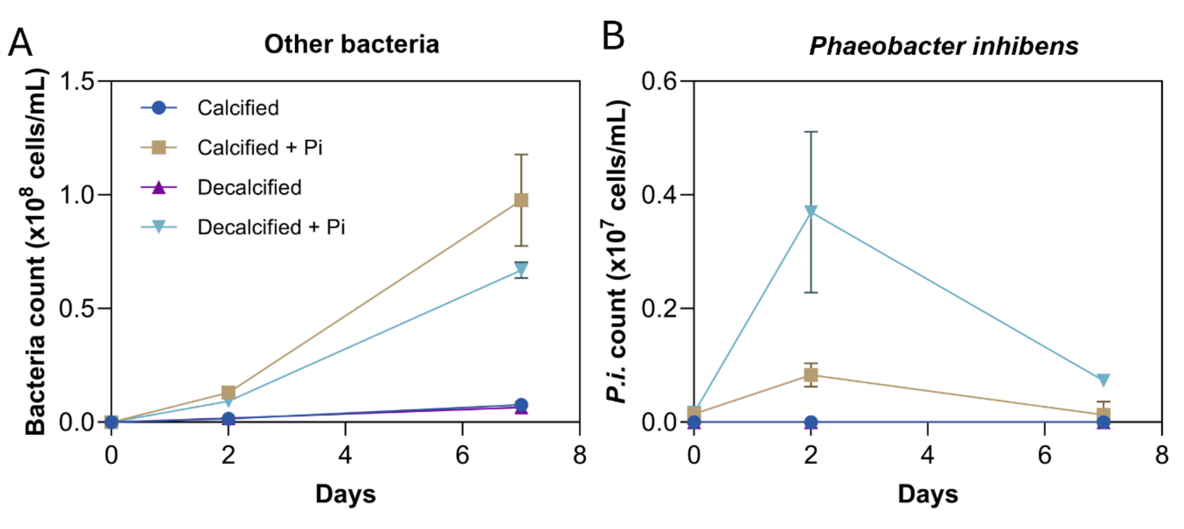


**Supplementary Figure 3. Counts of *Phaeobacter inhibens* and other bacteria when inoculated into calcified and decalcified *Coccolithus braarudii* cultures.** Bacteria counts as a function of time for (A) *P. inhibens,* monitored by plating samples on 100% marine broth agar plates and counting colony forming units (CFUs), for calcified *C. braarudii* cultures (blue circles), decalcified cultures (purple triangles), calcified cultures inoculated with *P. inhibens* (brown squares), and decalcified cultures inoculated with *P. inhibens* (inverted light blue triangles) and (B) all other bacteria in the algal culture.

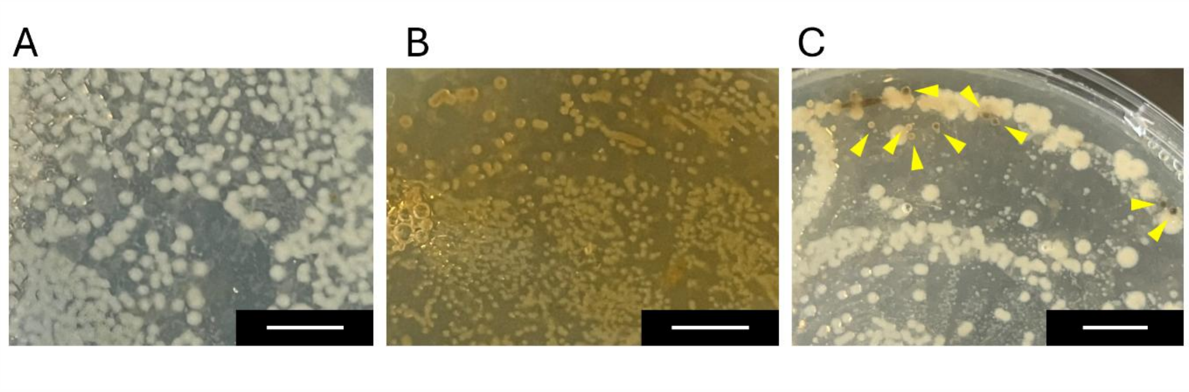


**Supplementary Figure 4. Bacterial colonies formed by communities from different *C. braarudii* treatments.** Colony forming units seen on 100% marine broth agar plates after 48 h incubation at 27 °C for (A) xenic *C. braarudii* culture (B) pure *P. inhibens* culture, (C) xenic *C. braarudii* culture inoculated with *P. inhibens* (*P. inhibens* colonies indicated with yellow triangles). Scale bars = 0.5 cm.

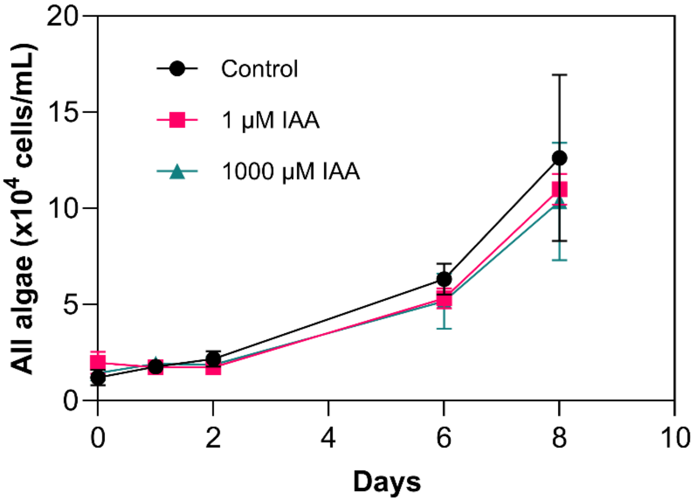


**Supplementary Figure 5. Growth of calcified *C. braarudii* cultures upon addition of indole-3-acetic acid (IAA).** Cell concentrations obtained by flow cytometry as a function of time for an untreated calcified *C. braarudii* culture (black circles) and for calcified cultures treated with indole-3-acetic acid at 1 µM (pink squares) or 1000 µM (green triangles). Points show the mean with SE bars for $n$ = 3 biological replicates. Comparison of treatments, two-way ANOVA, $F_{2,6}$ = 0.81, $p$ = 0.48.

Supplementary Video S1. ***Coccolithus braarudii* recalcification and cell division.** Timelapse microscopy video (one frame per hour, total duration 100 h) showing formation of coccoliths and cell division in EDTA decalcified *C. braarudii* cells.
